# The club cell circadian clock regulates temporal patterns in leukocyte trafficking in chronic allergic airways disease

**DOI:** 10.1101/2025.08.06.668949

**Authors:** Jafar Cain, Amlan Chakraborty, Vishvangi Deugi, Karolina Krakowiak, David Bechtold, Julie E Gibbs, Hannah J Durrington

## Abstract

**Rationale:** Asthma displays temporally variable symptoms which worsen overnight, corresponding with a nocturnal increase in airway eosinophils. The molecular clock within the club cell of the bronchial epithelium is a key driver of lung rhythmic processes, however, it’s role in chronic allergic airways disease (AAD) is not known. Elucidating the role of the club cell clock in regulating rhythmic inflammation in AAD could lead to new therapeutic advances.

**Objectives:** To investigate the club cell molecular clock regulation of leukocyte trafficking in chronic AAD.

**Methods:** *ccsp*-*bmal1* KO mice (which lack a functional clock in club cells) and littermate control mice underwent a 5-week chronic house dust mite (HDM) model of AAD, following which leukocyte populations and cytokines from blood, lung and airway compartments were quantified in a 24-hour time-course.

Airway epithelial cells were cultured and transepithelial electrical resistance measured to explore circadian variability in barrier permeability and impact of pharmacological modulation of the clock.

**Main Results:** Leukocyte populations accumulate in the blood, lung and airways of HDM exposed mice in a time-of-day dependent manner, with time of peak accumulation dependent on cell type. This temporal gating of leukocyte accumulation is controlled and coordinated by the club cell circadian clock, which also regulates airway barrier integrity. Targeting REVERBa (a component of the molecular circadian clock), was effective at modifying airway barrier permeability achieving reduced transepithelial leukocyte migration.

**Conclusions:** The club cell clock gates leukocyte trafficking signals and airway barrier integrity by time of day in chronic allergic airway inflammation.

## Introduction

Asthma is a chronic inflammatory disease of the lungs affecting over 350 million worldwide [1], commonly characterised by type-2 allergic airway inflammation, with T-helper (TH)2 cells and eosinophils being key drivers of disease. Many asthma symptoms exhibit temporal variations, typically being aggravated in the early hours of the morning [2]; implicating a role for the circadian clock in driving rhythmicity in allergic asthma. Indeed, populations who experience circadian disruption, such as through night shiftwork, experience a heightened prevalence and severity of asthma [3]. We previously found that the eosinophil chemoattractant eotaxin-1 peaks in concentration at 04:00 in sputum samples from patients with asthma [4], coinciding with physiological nadirs in lung function [4–6]. Moreover, it has been found that asthma exacerbations more commonly occur and are more likely to result in death at 04:00 [2, 7]. Although rhythmicity in asthma severity is well recognised, little is known about how the allergic inflammatory milieu of the lung changes over the course of the 24h day. Gaining mechanistic insight into drivers of rhythmic inflammation offers the opportunity to develop new therapeutic approaches.

Most mammalian cells have a molecular circadian clock which produces rhythmic gene outputs in 24-hour cycles. This molecular clock is centred on the actions of BMAL1 and CLOCK, transcription factors which drive the expression of clock-controlled genes [8, 9]. REVERBs and RORs are important clock outputs which form an accessory loop in the molecular circadian system and integrate the molecular clock with the immune system [10].

Airway epithelial cells (AECs), when damaged by inflammatory stimuli (including house dust mite (HDM) allergens), release alarmins which trigger a multifaceted immune response involving subsets of T cells (TH2s, TH9s) and eosinophils.

Eosinophils are key drivers of the pathophysiology in asthma. They enter the blood from the bone marrow and travel to the lung, releasing IL13, which induces goblet cell metaplasia and mucus hypersecretion [11, 12]. Moreover, IL13 damages AECs weakening the structural integrity of the epithelial barrier [13–15].

Club cells (CCs) have previously been identified as a population within the airway epithelium acting as local pacemakers [16, 17], coordinating and synchronising the actions of a wide range of local cell types [18–22]. Disrupting the CC circadian clock, via cell specific (CCSP^icre^) *Bmal1* ablation, leads to a loss of time-of-day variations in type-1 inflammatory lung disease models, and exacerbates disease severity [16, 17], however the role of the CC clock in coordinating rhythmicity in asthma remains to be explored.

Utilising an established mouse model of chronic allergic airways disease (AAD, reminiscent of asthma), here we show that the CC circadian clock is responsible for coordinating the time-of-day dependent accumulation of leukocytes in the lung and airways of mice. Furthermore, we demonstrate that the CC circadian clock is vital for maintaining physiological time-of-day dependent variations in airway barrier integrity and show that AAD-associated loss of barrier integrity can be restored through modulating the circadian clock in the airway epithelium.

## Methods

### Animals

Mice were maintained in the University of Manchester Biological Services Facility. All procedures were approved by the University of Manchester Animal Welfare and Ethical Review Body and carried out according to the Animals (Scientific Procedures) Act 1986. Mice were group housed under a 12 h:□12 h light/dark cycle with *ad libitum* access to food and water. C57BL6 mice were purchased from ENVIGO (*Huntingdon, United Kingdom*). CCSP-BMAL1 KO mice were bred in house as described elsewhere [16].

To induce chronic AAD female mice were dosed with HDM (Citeq; batches 15J02/20A05) [23] or vehicle (phosphate buffered saline (PBS)) three times per week for a period of 5 weeks. For further details see online supplement. At the end of the protocol, blood, broncho-alveolar fluid (BALF), lung and spleen tissue were collected. For *in-vivo* SR9009 dosing mice were intraperitoneally administered 100mg/kg SR9009 or an equivalent volume of DMSO every day at ZT6 for the final 14 days of the AAD protocol.

### EdU Dosing

100µl of 5-ethynyl-2’-deoxyuridine (EdU, 1mg/ml) (*ThermoFisher, USA)* was injected intraperitoneally to identify recently divided cells. EdU was detected by flow cytometry using the Click-iT EdU detection kit (*ThermoFisher, USA)* according to the manufacturer’s instructions.

### Tissue preparations for flow cytometry

Lung, BALF, and spleen samples were processed as previously described [16, 17, 24]. Blood samples were collected in EDTA coated tubes and processed according to an established protocol [25].

### Flow Cytometry Staining

Cells were stained for flow cytometric analysis according to established protocols [25, 26], (see online supplement). Analysis was carried out on a BD Fortessa flow cytometer.

### Total IgE ELISA

An ELISA (*Mouse IgE ELISA kit EMIGHE; ThermoFisher*, *USA)* was performed to analyse total serum IgE according to the manufacturer’s instructions.

### Bioplex Assay

BALF was analysed using Bio-Plex Pro ™ Mouse Chemokine Panel 23-Plex (Bio-Rad, #M60009RDPD) as per the manufacturer’s instructions.

### RT-qPCR

RNA was isolated from lung tissue samples, and RT-qPCR performed as previously described [23]. Primer sequences used can be found in Supplementary Table 1. Data from qPCR assays were normalised against the housekeeping gene *Rn18s*.

### Immunoblotting

Protein was isolated and quantified from lung tissue and immunoblotting performed as described previously with antibody and concentrations details in Supplementary Table 4. [27, 28]

### Histology

Established protocols were used for lung tissue histological and immunohistochemical staining [29, 30].

### Air-Liquid Interface Cell Culture

Tracheal epithelial cells from CCSP-BMAL1 KO and wild-type male mice (10-13 weeks) were cultured as previously described [31]. Cells were synchronised with dexamethasone (200ng/ml, 2h) before transepithelial electrical resistance (TEER) measurements every 4h for 96h using an electrovoltmeter, in the presence/absence of IL13 (20ng/ml) and/or SR9009 (5µM) or DMSO control.

### Statistical Analysis

Statistical analysis was performed using GraphPad Prism software version 10. Two-way, and Three-way ANOVAs were used to determine significance of variables, with post-hoc Sidak’s/ Dunnett’s multiple comparison tests as appropriate. A p-value ≤ 0.05 was considered statistically significant. Cosinor analysis was conducted in DiscoRhythm v1.2.1 [32].

## Results

### Leukocyte populations vary by time-of-day in chronic allergic airways disease (AAD)

Using a chronic model of AAD, we sought to establish time-of-day dependent patterns in leukocyte accumulation in the blood, BALF, lungs, and spleen. Mice were sampled at Zeitgeber Time (ZT) 0 (time of lights on and start of the rest phase), ZT6 (mid-rest phase), ZT12 (lights off and start of the active phase) and ZT18 (mid-active phase). After 5 weeks of sensitisation with HDM (batch 15J02) (**Fig1A**), circulating blood eosinophils were quantified by flow cytometry (Siglec-F+, CD11b+) in HDM-exposed mice and PBS-treated controls over the 40-hours following the final exposure (**Fig 1B**). Under normal conditions, circulating eosinophils were elevated at ZT6 compared to all other time points, a pattern significantly amplified in the presence of AAD (**Fig1B**). EdU dosing 6 hours prior to sample collection revealed that the overwhelming majority (∼95%) of eosinophils isolated at ZT6 – from both PBS and HDM treated mice - were EdU+ and thus newly divided cells (**Fig1C**).

**Figure 1:**
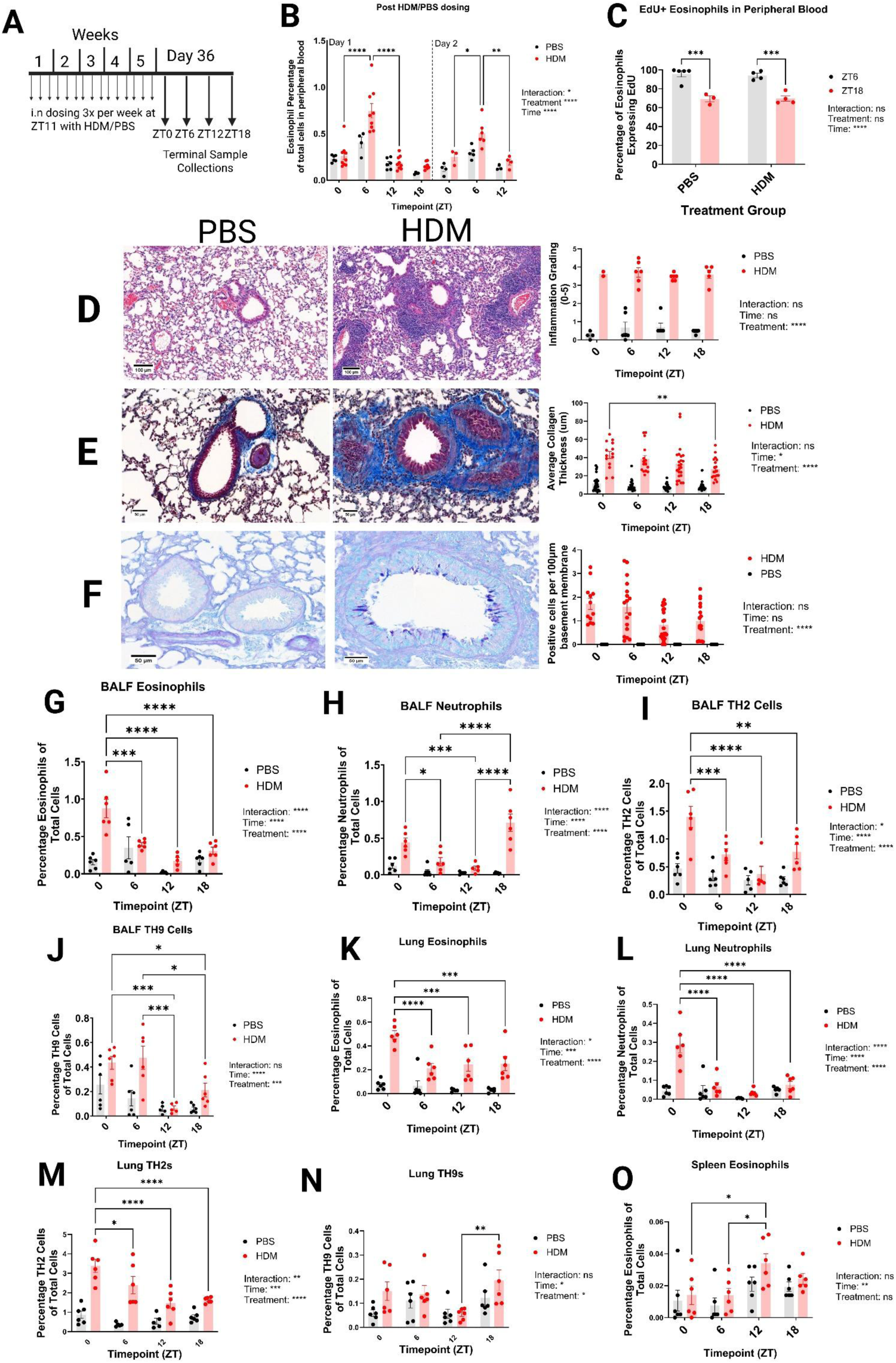
Leukocyte populations vary by time-of-day in chronic allergic airways disease (AAD) (A) – Schematic overview of 5-week dosing protocol and sampling timepoints. (B) – Quantification of eosinophils in peripheral blood samples across 7 consecutive 6-hourly timepoints following the final dose of HDM/PBS. (C) – Percentages of EdU positive eosinophils sampled at ZT6 and ZT18 in the first day post-final HDM/PBS dose administration. (D) - Representative images of haematoxylin and eosin-stained lung sections from PBS/HDM challenged mice on day 36 of the chronic AAD protocol and quantification of inflammation scoring across all 6-hourly timepoints. (E) - Representative images of Masson’s Trichrome-stained lung sections from PBS/HDM challenged mice on day 36 of the chronic AAD protocol and quantification of peribronchiolar collagen deposition across all 6-hourly timepoints. (F) - Representative images of alcian blue periodic acid schiff stained lung sections from PBS/HDM challenged mice on day 36 of the chronic AAD protocol and quantification of positive stained cells per 100µm of basement membrane across all 6-hourly timepoints. (G – O) – Quantification of immune cell subsets in BALF, and Lung sampled across 4 6-hourly timepoints on day 36 of the chronic HDM exposure protocol. (B-O) – all data presented as mean +/- SEM analysed by two-way ANOVA with post-hoc Šidak’s multiple comparison tests. N= 3-9. *P< 0.05, **P< 0.01, ***P< 0.001, ****P< 0.0001. First level output of two-way ANOVA shown as Interaction / Time / Treatment with corresponding P value indicated.

Meanwhile, at ZT18 only approximately 70% of circulating eosinophils were EdU+ and so newly divided, suggesting that at this later timepoint eosinophils may recirculate more readily (**Fig1C**). We observed the same time-of-day oscillations in both PBS and HDM-treated mice, suggesting that this rhythmicity in circulating eosinophils is present under homeostatic conditions, and then amplified in the presence of AAD (**Fig1B-C**).

In the lung, chronic HDM exposure induced gross pathological changes which were stable across timepoints (**Fig1D-F**). Both haematoxylin and eosin (**Fig1D**), and Alcian Blue Periodic Acid Schiff (**Fig1F**) staining revealed treatment-dependent inflammatory changes in the lung tissue however this did not vary significantly by time-of-day. Due to the chronic nature of the protocol, we did not anticipate finding significant time-dependent differences in peribronchiolar collagen deposition, although analysis suggested that in ZT0 samples there was more deposition compared to ZT18 samples (**Fig1E**).

Chronic HDM exposure precipitated both an absolute and a relative increase in eosinophil, neutrophil, TH2, and TH9 cell populations in the airways (measured in BALF), especially during the rest phase (ZT0, ZT6) (**Fig1 G –J; Sup Fig1 A - F)**. Likewise, in the lung, there was a strong treatment effect associated with HDM exposure, with greater populations of eosinophils and TH2 cells occurring during the rest period (**Fig1 K - N**), however this was not found to apply to Treg cells **(Sup Fig1 G)**. Additionally, splenic eosinophil populations in both PBS and HDM-treated mice were elevated at ZT12 (**Fig1O**).

Overall, our findings show that despite all HDM-exposed mice reaching comparable levels of gross pathology, distinct compartment specific time-of-day patterns in leukocyte populations exist. Inflammatory cell populations in the lung and BALF peak at ZT0, and trough at ZT12. In the blood, eosinophils exhibit daily oscillatory cycles, being most abundant at ZT6, a timepoint associated with increased eosinophilopoiesis.

### Airway cytokines vary by time-of-day in allergic airways disease

Having observed daily variation in accumulation of pro-inflammatory leukocytes within the allergically inflamed lung we next examined whether cytokine signals associated with these cells exhibited time-of-day variation. We find that in HDM (batch 15J02)-exposed mice significant time-of-day dependent variations exist in a range of BALF cytokines (**Fig2A**).

Whilst TH2 cells peak in the BALF and lung of HDM-exposed mice at ZT0 **(Fig1I, M)**, TH2 cytokines, namely IL4, IL13 and IL5, were most elevated at ZT6 **(Fig2 B - D)**.

Eotaxin-1, a key eosinophil chemoattractant, peaked at ZT18 **(Fig2J)**, 6-hours preceding the peak in BALF eosinophil populations **(Fig1G)**. IL9 levels were significantly raised during the mouse active phase (ZT12/18) (**Fig2D**), a finding that correlates with patterns observed in TH9 cell numbers in the BALF (**Fig1J**).

Whilst some elevation in type-1 cytokines was expected in this model of AAD **(Supp Fig1 K - M)**, only IL12, predominantly produced by antigen presenting cells, exhibited temporal variation, being most elevated at ZT12 in the BALF of HDM-exposed mice **(Fig2F)**.

Leukopoeitic cytokines were found to be elevated in BALF samples from HDM-exposed mice compared to PBS controls and significant time-of-day dependent variations were also observed in the concentration of GM-CSF, IL5, and GCSF **(Fig2I - K)**. GCSF and GM-CSF were found to peak in concentration during the mouse active phase **(Fig2K, J)**, whilst IL5 peaked at ZT6 in the BALF of HDM-exposed mice (**Fig2I**) Overall, these findings reveal that in the allergically inflamed lung, distinct temporal rhythms in cytokine production exist varying according to cytokine type.

### In AAD the club cell circadian clock regulates time-of-day changes in immune cell populations within the lung and airway

Building on our observations that HDM-exposure was associated with time-of-day dependent patterns of immune cell accumulation in the BALF, we repeated the chronic HDM model (here with HDM batch 20A05) in mice lacking a functional circadian clock in the club cells of the airway epithelium, CCSP-BMAL1 KO mice **(Fig3A)**.

Histological analyses of lung tissue assessed mucin production, immune cell infiltration, and peribronchiolar collagen deposition (**Fig3B – D)**. While no time-of-day variations were found in mucin staining in WT HDM-exposed mice, CCSP-BMAL1 KO mice showed increased mucin at ZT12 compared to ZT0 (**Fig3B**).

We noted that IgE concentrations in the serum varied by time-of-day, being significantly elevated at ZT12 compared to ZT0 in WT mice (Bmal^flox/flox^), but not in HDM-exposed CCSP-BMAL1 KO mice where IgE was significantly reduced (**Supp Fig2J**). B-cell numbers in peripheral blood, did not vary by time-of-day in WT and CCSP-BMAL1 KO mice **(Supp Fig2K)** suggesting that the variance in IgE concentration is linked to functional activity rather than increases in the size of the B-cell population.

There was an increase in BALF total cell numbers after HDM-exposure in both WT and CCSP-BMAL1 KO mice, which did not vary between timepoints **(Supp Fig 2A**). The composition of BALF samples however differed significantly between genotypes and timepoints. In WT mice, both eosinophils and TH2 cell populations were raised at ZT0 compared to ZT12 (as found previously, **Fig1G, I**), a phenomenon that was lost in CCSP-BMAL1 KO mice **(Fig3E, H)**. Furthermore, we found that the CCSP-BMAL1 KO genotype was associated with increased BALF neutrophilia, in response to HDM-exposure, compared to WT mice, regardless of time **(Fig3F)**. We noted also that the CCSP-BMAL1 KO genotype was associated with increased TH9 cell populations in both the lung and BALF (**Fig3F, J**). TH9 cell populations were most elevated at ZT12 in HDM-exposed CCSP-BMAL1 KO mice (**Fig3F, J**), something which was not seen in WT HDM-exposed mice in this experiment (**Fig3F, J**), nor in previous experiments with WT mice, where ZT0 and ZT6 had been associated with increases in TH9 cell populations **(Fig1J)**.

**Figure 2:**
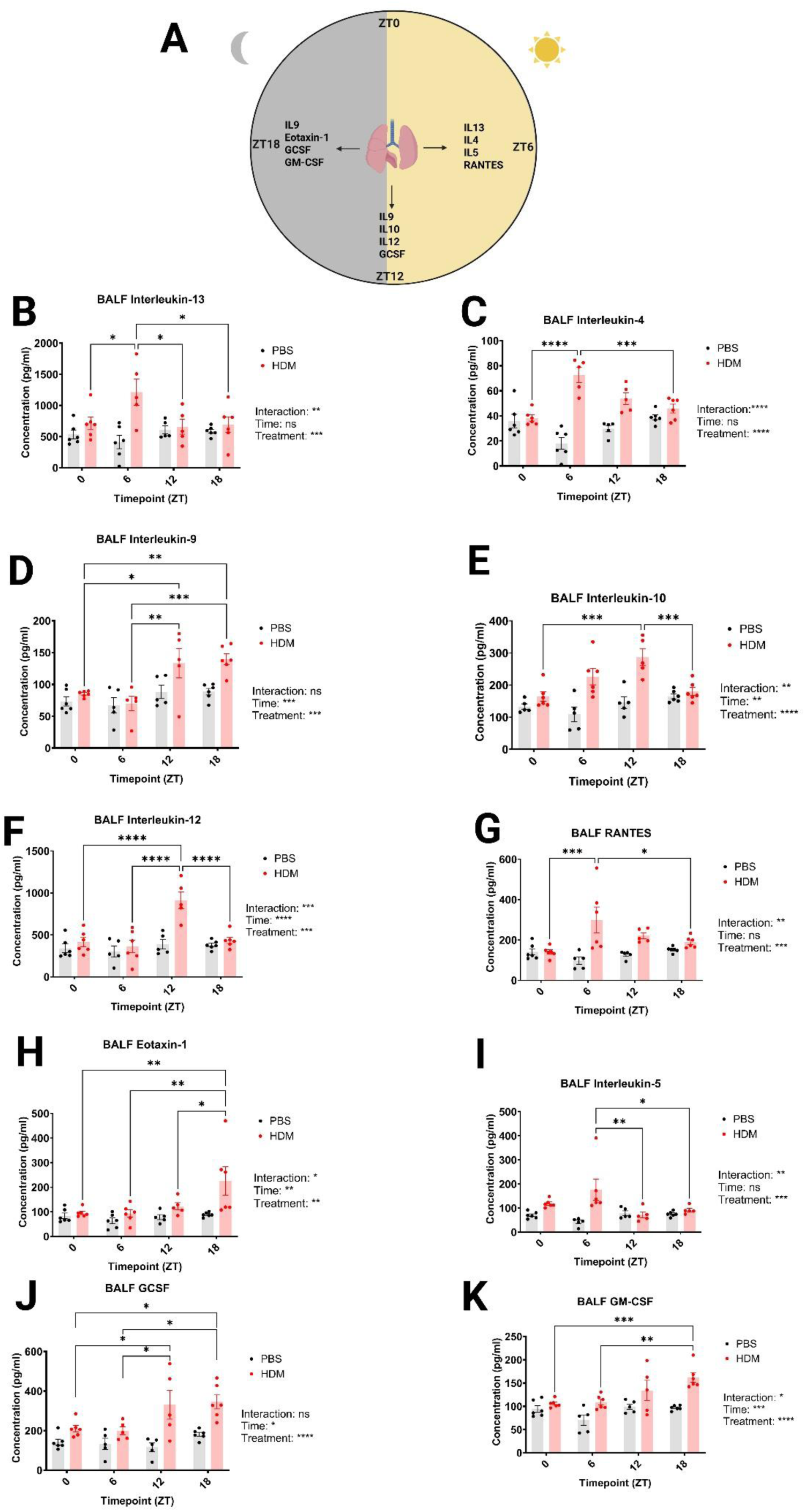
BALF cytokine concentrations vary by time-of-day in the HDM model of allergic airways disease (A) – Schematic outline of the time-of-day at which important BALF cytokines are significantly elevated. At ZT6, cytokines such as IL13, IL4, IL5, and RANTES, associated with TH-cell activity peak in the BALF of HDM-exposed mice. At ZT12 IL9, IL10, and IL12 are found to peak in their respective concentrations in the BALF of HDM-exposed mice. Eotaxin-1, GCSF, and GM-CSF are found to reach their peak concentrations in the BALF at ZT18, a timepoint in which IL9 is also elevated. (B-K) - Quantification of cytokines present in BALF samples of HDM/PBS treated mice taken at 4 6-hourly intervals on day 36 of the chronic HDM exposure protocol. (B-K) - all data presented as mean +/- SEM analysed by two-way ANOVA with post-hoc Šidak’s multiple comparison tests. N= 5-6. *P< 0.05, **P< 0.01, ***P< 0.001, ****P< 0.0001. First level output of two-way ANOVA shown as Interaction / Time / Treatment with corresponding P value indicated.

Flow cytometric analysis of lung samples revealed that eosinophils, along with TH2 cells, were significantly raised in HDM-exposed WT mice at ZT0, compared to ZT12 (**Fig3I, J**). This temporal variation was lost in samples from CCSP-BMAL1 KO mice (**Fig 3I, J)**. Further examination of eosinophil populations revealed that this time-of-day difference was driven by changes in CD11c^mid^ (inflammatory [33]) eosinophils (**Fig3I**) rather than a baseline expansion of CD11c^low^ (tissue resident [33]) eosinophils **(Sup Fig2 F)**. Curiously, within the lung we again observed that the CCSP-BMAL1 KO genotype, regardless of time, was associated with increases in TH9 cell populations, compared to WT littermate controls **(Fig3K)**.

**Figure 3:**
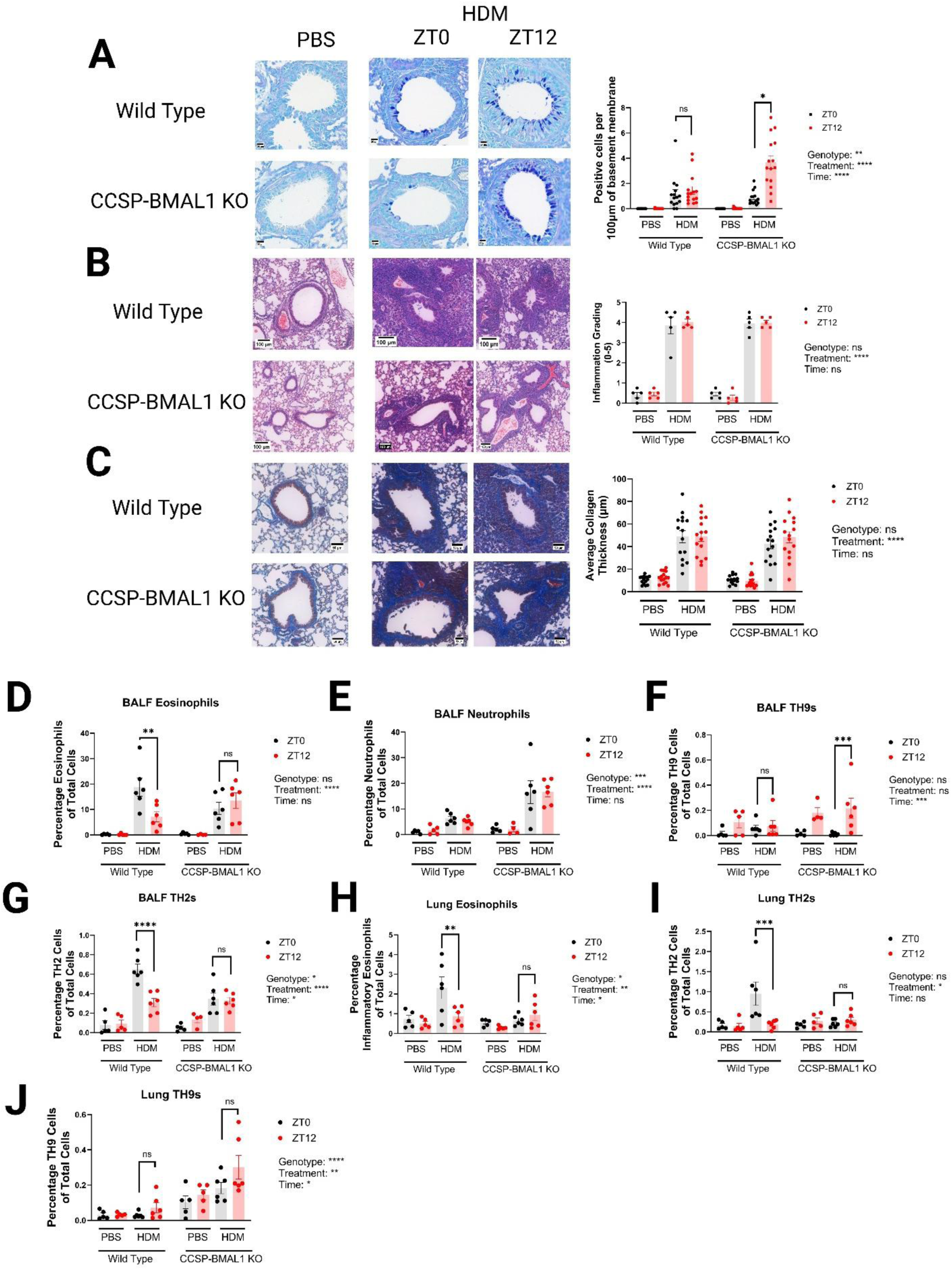
The club cell circadian clock is responsible for gating time-of-day differences in immune cell accumulation in the lung and BALF of HDM-exposed mice (A) – Representative images and quantification of positive mucin staining in AB-PAS stained FFPE lung sections taken at ZT0 and ZT12 from HDM/PBS treated CCSP-BMAL1 KO mice and wild type littermate controls, shown as number of positive cells per 100µm of basement membrane. (B) – Representative images and quantification of H&E stained FFPE lung sections, taken at ZT0 and ZT12 from HDM/PBS treated CCSP-BMAL1 KO mice and wild type littermate controls, according to the scale detailed in Denny et al 2015. (C) – Representative images and quantification of the extent of peribronchiolar collagen deposition in FFPE lung sections, taken at ZT0 and ZT12 from HDM/PBS treated CCSP-BMAL1 KO mice and wild type littermate controls, measured in Aperio Case Viewer. (E – K) – Quantification of immune cell subsets in BALF and Lung of HDM/PBS treated CCSP-BMAL1 KO mice and wildtype littermate controls, sampled at ZT0 and ZT12 on day 36 of the chronic HDM exposure protocol. (A-K) – all data presented as mean +/- SEM analysed by three-way ANOVA with post-hoc Šidak’s multiple comparison tests. N= 5-16. *P< 0.05, **P< 0.01, ***P< 0.001, ****P< 0.0001. First level output of three-way ANOVA shown as Genotype / Treatment / Time with corresponding P value indicated.

### A functioning club cell circadian clock is needed to maintain time-of-day dependent variations in several BALF cytokines

Quantification of cytokines within the BALF following HDM (batch 20A05) exposure revealed an overall, genotype-associated increase in IL9 in CCSP-BMAL1 KO mice, regardless of time **(Fig4B)**. This aligns with our observation that TH9 cell populations were increased in the lungs and BALF of these same mice **(Fig3G, H)**. IL5, a key marker of type-2 inflammation, was also elevated in HDM-exposed WT mice at ZT0 compared to ZT12, a finding not observed in HDM-exposed CCSP-BMAL1 KO mice (**Fig4A)**.

**Figure 4:**
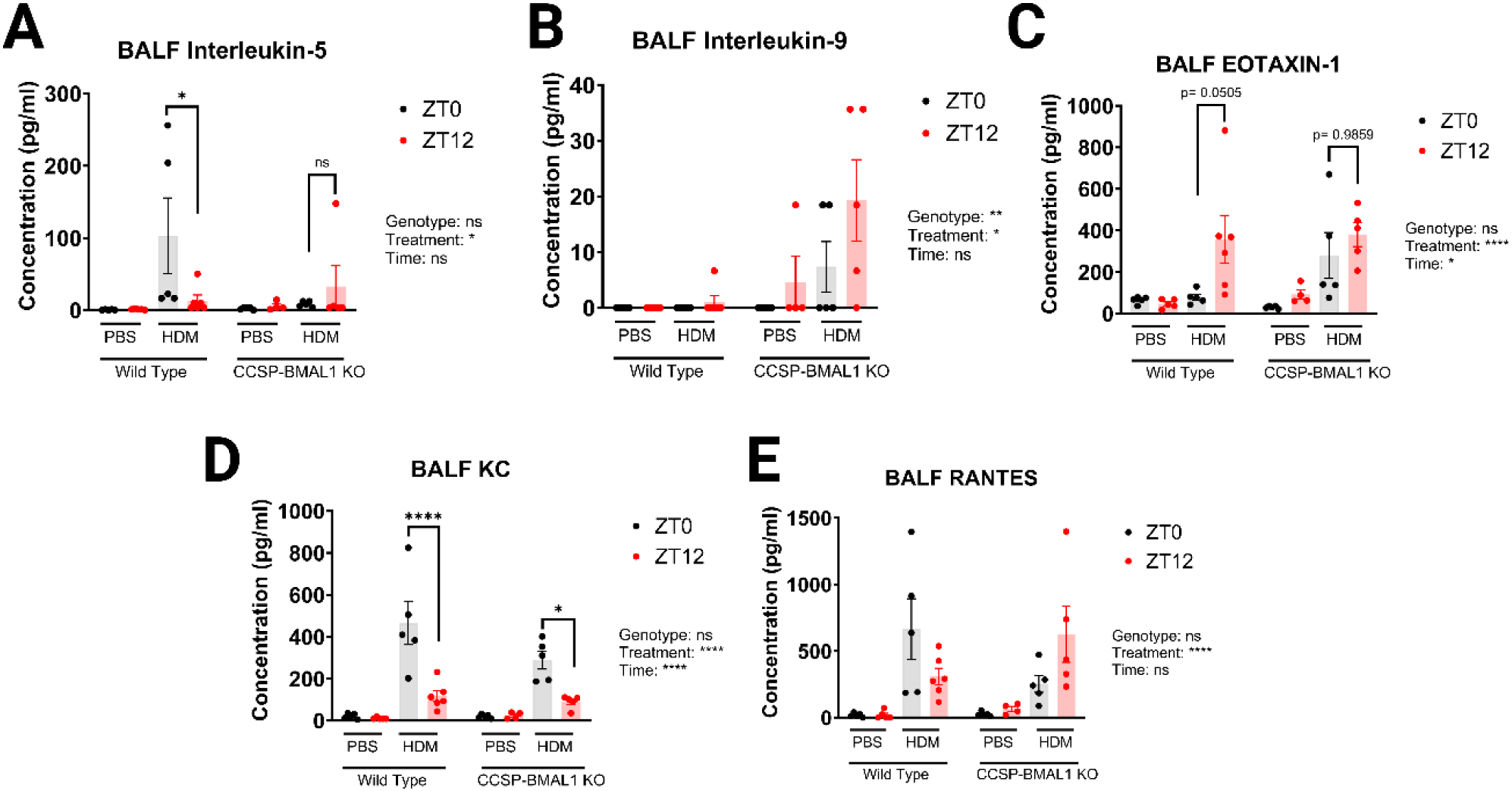
A functioning club cell circadian clock is needed to maintain time-of-day dependent variations in a number of BALF cytokines (A-E) – Quantification of cytokines present in BALF samples of HDM/PBS treated CCSP-BMAL1 KO mice and wild type littermate controls taken at ZT0 and ZT12 on day 36 of the chronic HDM exposure protocol. All data presented as mean +/- SEM analysed by three-way ANOVA with post-hoc Šidak’s multiple comparison tests. N= 5-6. *P< 0.05, **P< 0.01, ***P< 0.001, ****P< 0.0001. First level output of three-way ANOVA shown as Genotype / Time / Treatment with corresponding P value indicated.

Crucially, eotaxin-1 concentrations did exhibit time-of-day variation according to the three-way ANOVA, however post hoc comparisons demonstrated that this temporal variation was just outside the threshold for significance (p= 0.0505) in WT HDM-exposed mice **(Fig4C)**. HDM-exposed CCSP-BMAL1 KO mice however did not show any time-of-day variation in eotaxin-1 **(Fig4C)**.

### The club cell circadian clock is essential for the temporal regulation of airway barrier integrity

Based on our observations that leukocyte populations in the BALF exhibit time-of-day variations, we sought to assess whether this could be attributed to variability in epithelial tight junction (TJ) function and if this could further be related to the CC circadian clock. Immunohistochemical investigation of WT mice revealed that TJ proteins ZO.1 and Occludin were most abundant in PBS-treated mice at ZT6, and expression was reduced following HDM exposure (**Supp Fig4 A - J**). Through RT-qPCR analysis of lung tissue from naïve mice at ZT6 and ZT18, we found that gene expression of *Zo.1* and *Ocln* varies by time-of-day in WT mice **(Fig5A, B)** but this variation was strikingly lost when *Bmal1* was ablated from CCs **(Fig5A, B)**. Building on this finding we isolated primary tracheal epithelial cells and differentiated them into a monolayer of mature epithelial cells using an Air Liquid Interface (ALI) protocol **(Fig5C)**. We then tested the rhythmicity (using cosinor analysis) of the transepithelial electrical resistance (TEER; a measure of TJ integrity) across ALI cell membranes grown from WT and CCSP-BMAL1 KO derived cells **(Fig5D)**. Following synchronization with dexamethasone (dex), cosinor analysis revealed significant rhythmicity in the TEER of WT ALI cell cultures **(Fig5D)**. In contrast, ALI cultures from CCSP-BMAL1 KO mice did not exhibit temporal variation in TEER measurements **(Fig5D)**.

Dosing ALI cultures with SR9009, a synthetic REVERBa ligand, increased TEER in both WT and CCSP-BMAL1KO groups, suggesting increased barrier integrity (**Fig5E-H**). Importantly, we note that whilst *Bmal1* ablation renders the circadian clock in the club cell non-functional, residual levels of REVERBa are still found in these cells[17], therefore we postulated that sufficient protein existed for the REVERBa ligand SR9009 to exert effect. Following SR9009 treatment, cells were exposed to IL13, a cytokine associated with allergic inflammation and with barrier disruption in particular [34, 35]. Whilst IL13 exposure did reduce the TEER values in both WT and CCSP-BMAL1 KO cultures, this was not by enough to reduce barrier integrity below baseline (**Fig5E-H**). Indeed, cultures exposed to IL13, in the absence of SR9009 pre-treatment, exhibited a significant reduction in TEER below their original baseline values (**Fig5E-H**). Reversing the order of treatments further demonstrated that dosing epithelial cells with SR9009, post-IL13 exposure, restored TEER values to baseline **(Supp Fig5A-D)**.

Since HDM exposure is associated with reduced TJ gene expression *in-vivo* (**Supp Fig5 E, F**), we hypothesised based on our *in-*vitro experiments that treating HDM-exposed mice with SR9009 could help restore barrier integrity. Analysis of *Zo.1* and *Ocln* gene and protein expression revealed that SR9009 treatment was indeed able to increase the expression of these tight junction components in the lung tissue of mice exposed to HDM (**Fig 5I, J, K**).

**Figure 5:**
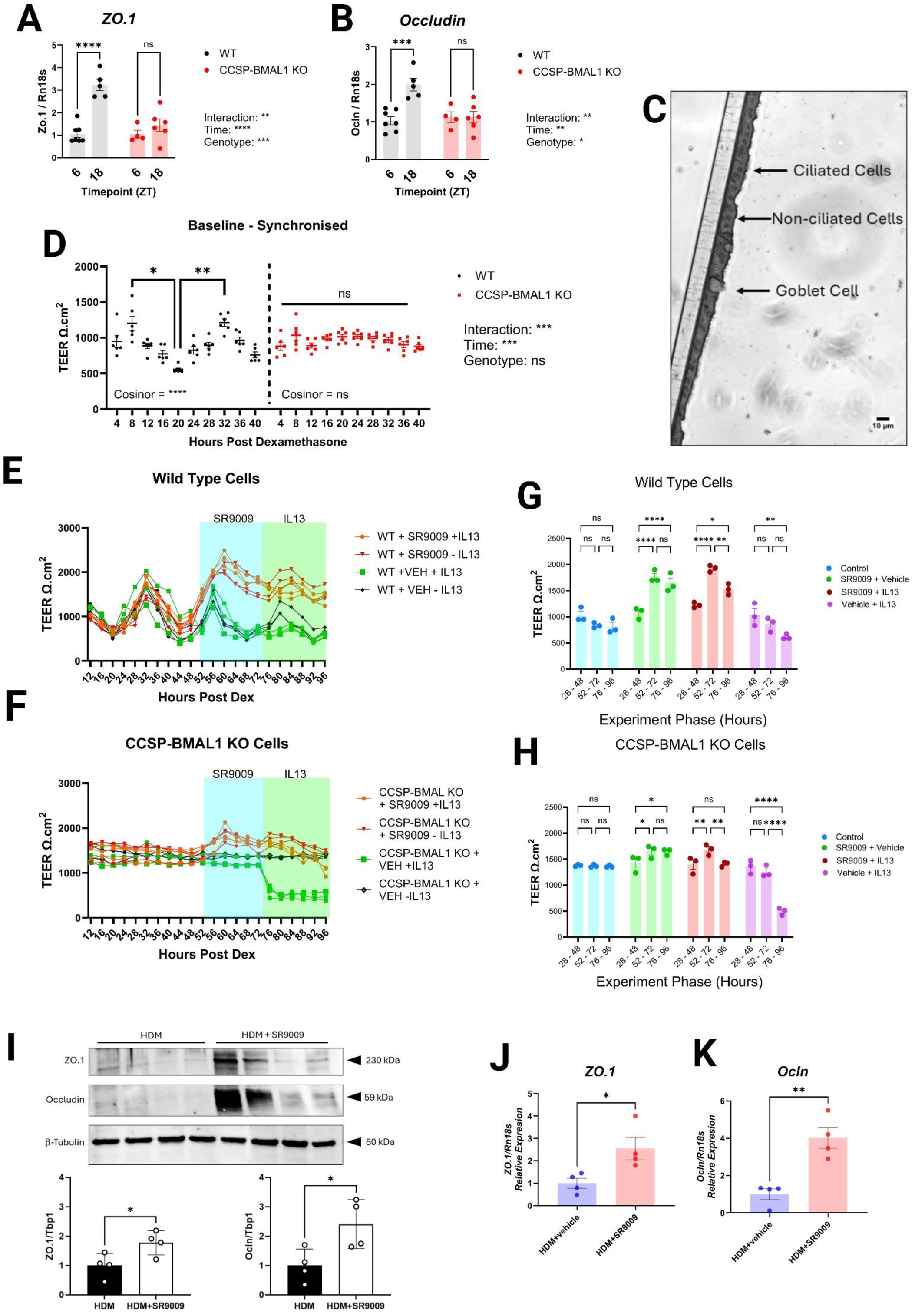
The club cell circadian clock is essential for the temporal regulation of airway barrier integrity (A) – qPCR data showing the relative expression of tight junction gene *Zo.1* at ZT6 and ZT18 in whole lung samples taken from male CCSP-BMAL1 KO mice and wild type littermate controls. (B) - qPCR data showing the relative expression of tight junction gene *Ocln* at ZT6 and ZT18 in whole lung samples taken from male CCSP-BMAL1 KO mice and wild type littermate controls. (A-B) –qPCR data was normalised to the wild type ZT6 group, and relative to the expression of housekeeping gene *Rn18s,* shown as mean +/- SEM, analysed by two-way ANOVA with post-hoc Sidak’s multiple comparison tests, n= 5-6, *P< 0.05, **P< 0.01, ***P< 0.001, ****P< 0.0001. First level output of two-way ANOVA shown as Interaction / Time / Genotype with corresponding P value indicated. (C) – Representative image of an FFPE section of Air-Liquid interface cultured epithelial cells taken using a EVOS M5000 light microscope. (D) – Trans-epithelial Electrical Resistance measurements taken from wild type and CCSP-BMAL1 air-liquid interface airway epithelial cell cultures every 4 hours for 40 hours post-synchronisation of internal clocks using dexamethasone. Data shown as mean +/- SEM, n= 6 per timepoint per genotype, analysed by two-way repeated measurements ANOVA with post-hoc Tukey’s t-tests, *P< 0.05, **P< 0.01, ***P< 0.001, ****P< 0.0001. First level output of two-way ANOVA shown as Interaction / Genotype / Time with corresponding P value indicated. Rhythmicity was assessed using Cosinor analysis in DiscoRhythm, *P< 0.05, **P< 0.01, ***P< 0.001, ****P< 0.0001. (E-F) - Trans-epithelial Electrical Resistance measurements taken from wild type (E) and CCSP-BMAL1 KO (F) air-liquid interface airway epithelial cell cultures every 4 hours from 12 hours post-synchronisation of internal clocks using dexamethasone to 96 hours post-synchronisation. At hour 48 cultures were dosed with SR9009 or DMSO as vehicle control, before being subsequently exposed to IL13 at hour 72. N=3 per genotype per treatment group. (G – H) The average TEER for each experimental phase (hours 28 – 48, 52 – 72, 76 – 96) were calculated for each individual well. N=3 per treatment group per experiment phase, analysed by two-way ANOVA with post-hoc Tukey’s T-tests. Data shown as mean +/- SEM, *P< 0.05, **P< 0.01, ***P< 0.001, ****P< 0.0001. (I) – Western Blots were performed to assess the relative quantity of ZO.1 and Occludin proteins in lung tissue lysates taken from HDM exposed mice +/- SR9009 treatment. (J) - qPCR data showing the relative expression of tight junction gene *Zo.1* in lungs taken from HDM-exposed mice +/- treatment with SR9009. (K) - qPCR data showing the relative expression of tight junction gene *Ocln* in lungs taken from HDM-exposed mice +/- treatment with SR9009. (I-K) – n=4 per treatment group. Data shown as mean +/- SEM, analysed by unpaired two-tailed t-tests, *P< 0.05, **P< 0.01, ***P< 0.001, ****P< 0.0001.

## Discussion

This study reveals, for the first time, that immune cell accumulation in the lungs and airways of mice with chronic AAD follows a 24-hour rhythm. The club cell circadian clock plays a central role in coordinating this temporal variation. Additionally, we demonstrate that the club cell regulates airway barrier rhythmicity, likely via REVERBa, linking the circadian clock to barrier integrity.

Our clinical studies highlight sputum eosinophils as highly rhythmic in patients with severe asthma, prompting exploration of this cell type, key to the pathogenesis of allergic asthma, in our model [4]. Like we have found in humans, eosinophils accumulate in the lungs of mice during the rest-phase, demonstrating the relevance of this model for investigating human disease.

The CC has emerged as a crucial cell type responsible for gating time-of-day responses to allergen challenge in this study as well as similar responses to LPS and viral challenges [16, 17]. We suggest that the CC coordinates the time-of-day dependent changes in BALF eosinophils and TH-cell populations through regulating both the physical integrity of the airway barrier as well as the release of chemokines such as eotaxin-1 which attract key leukocytes to the inflamed airway [36, 37].

Clock-controlled regulation of TJ genes (*Zo.1, Ocl*) has been reported in other tissues, and our findings extend this to the lung, and specifically to CCs of the airway epithelial barrier [38, 39]. We, and others, have found that compounds that activate REVERBa such as SR9009, are able to reduce barrier permeability in ex-vivo experiments [40, 41]. The ability of SR9009 to reduce barrier permeability, and thereby potentially limit the influx of allergens and activation of immune pathways perpetuating airway barrier damage, raises the potential of utilising inhaled clock-targeting drugs as novel asthma therapy.

These findings have significant implications for our understanding of allergic asthma. We have previously shown that disease severity in HDM-exposed mice is linked to timing of allergen exposure [23], and results presented here suggest that this is likely due to changes in airway permeability over the 24h day. Administering allergens when the barrier is more permeable may exacerbate inflammation, highlighting the need to consider dosing time in experiments. Increased permeability also facilitates leukocyte migration, potentially explaining time-of-day immune variations in BALF and patient sputum [4, 23].

In line with findings here, time-of-day differences in CD4+ T-cell populations in allergen-challenged mice have been previously observed, but TH-cell subsets and the molecular clock’s role remained unclear [42]. We found that *Bmal1* ablation in club cells disrupted TH2 rhythmicity, while TH9 cells exhibited a phase shift and increased amplitude. This increase in TH9 cells in CCSP-BMAL1 KO mice was accompanied by elevated BALF IL9 levels. Further work to investigate the role of the CC clock mechanism in regulating TH9/IL9 pathways is warranted given the role of TH9 in asthma [43–46].

Additionally, we focussed specifically on how the CC molecular clock regulates immune cell trafficking in AAD. CCSP-BMAL1 KO mice exhibited loss of temporal variation in cellular composition of the BALF, with reduced inflammatory cell populations at ZT0 compared to WT littermates. Future work should elucidate how this translates to alterations in lung function and thus the extent to which the CC molecular clock controls daily rhythms in AHR.

In conclusion, we have shown that the CC coordinates time-of-day dependent changes in eosinophil and TH-cell populations through regulating both the physical integrity of the airway barrier, and the release of chemokines. These findings have multiple clinical implications, emphasising the need to take time into consideration when diagnosing, assessing and treating allergic asthma. Furthermore, these data highlight the potential for clock-modulatory compounds as potential therapeutics in AAD.

---

**Supplementary Figure 1.**
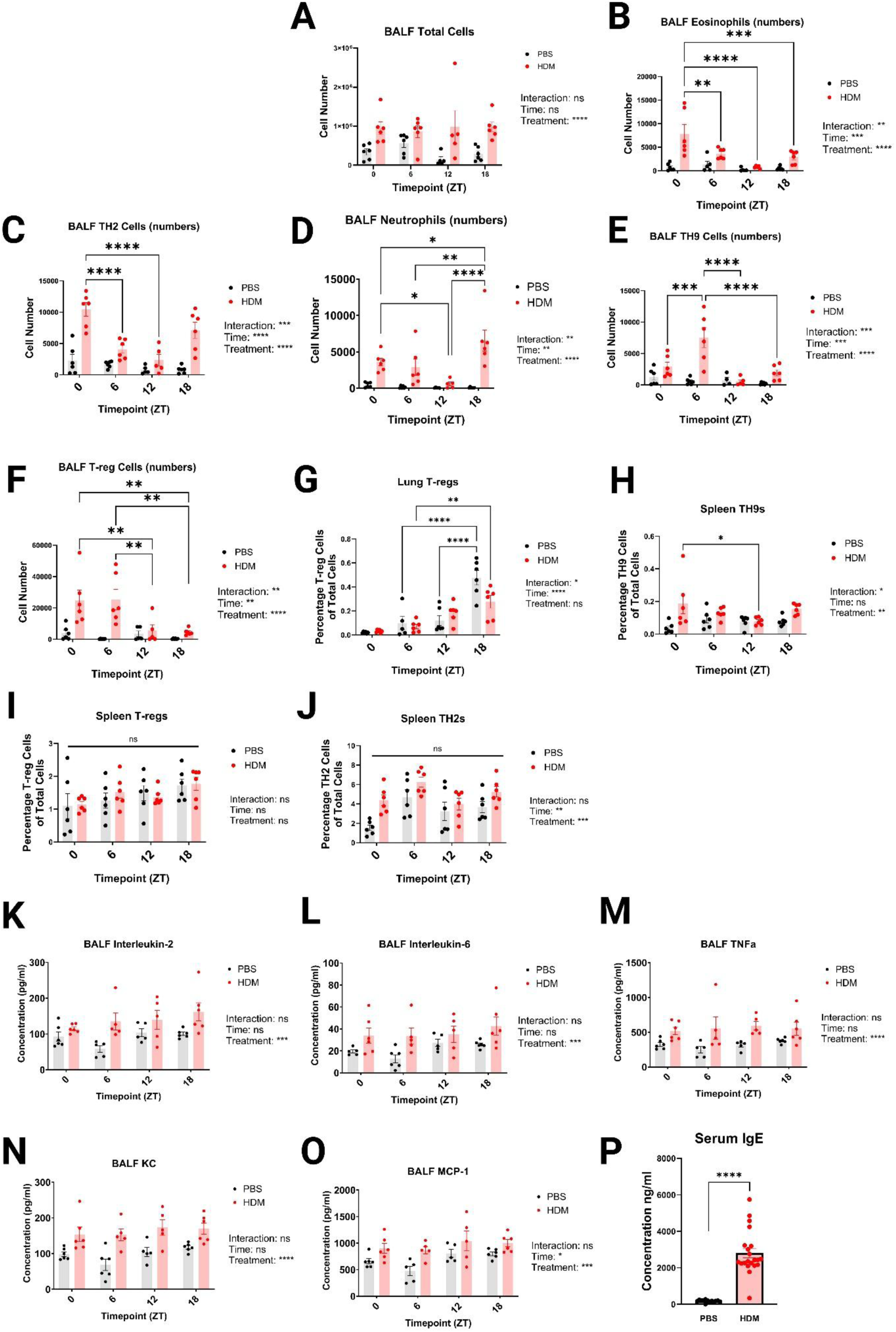
(A-J) – Immune cell subsets from HDM/PBS treated mice were quantified by flow cytometry from BALF, lung and spleen samples taken on day 36 of the chronic HDM exposure protocol at 4 6-hourly intervals. (K – O) Quantification of cytokines (IL2, IL6, TNFa, KC, and MCP-1) in BALF samples taken on day 36 of the chronic HDM exposure protocol at 4 6-hourly intervals. (P) – Serum IgE was quantified by ELISA from peripheral blood samples taken from HDM/PBS treated mice on day 36 of the chronic HDM exposure protocol. (A-P) – All data presented as mean +/- SEM, analysed by one/two-way ANOVA as appropriate with post-hoc Sidak’s multiple comparison tests. N= 5 – 21, *P< 0.05, **P< 0.01, ***P< 0.001, ****P< 0.0001.

**Supplementary Figure 2.**
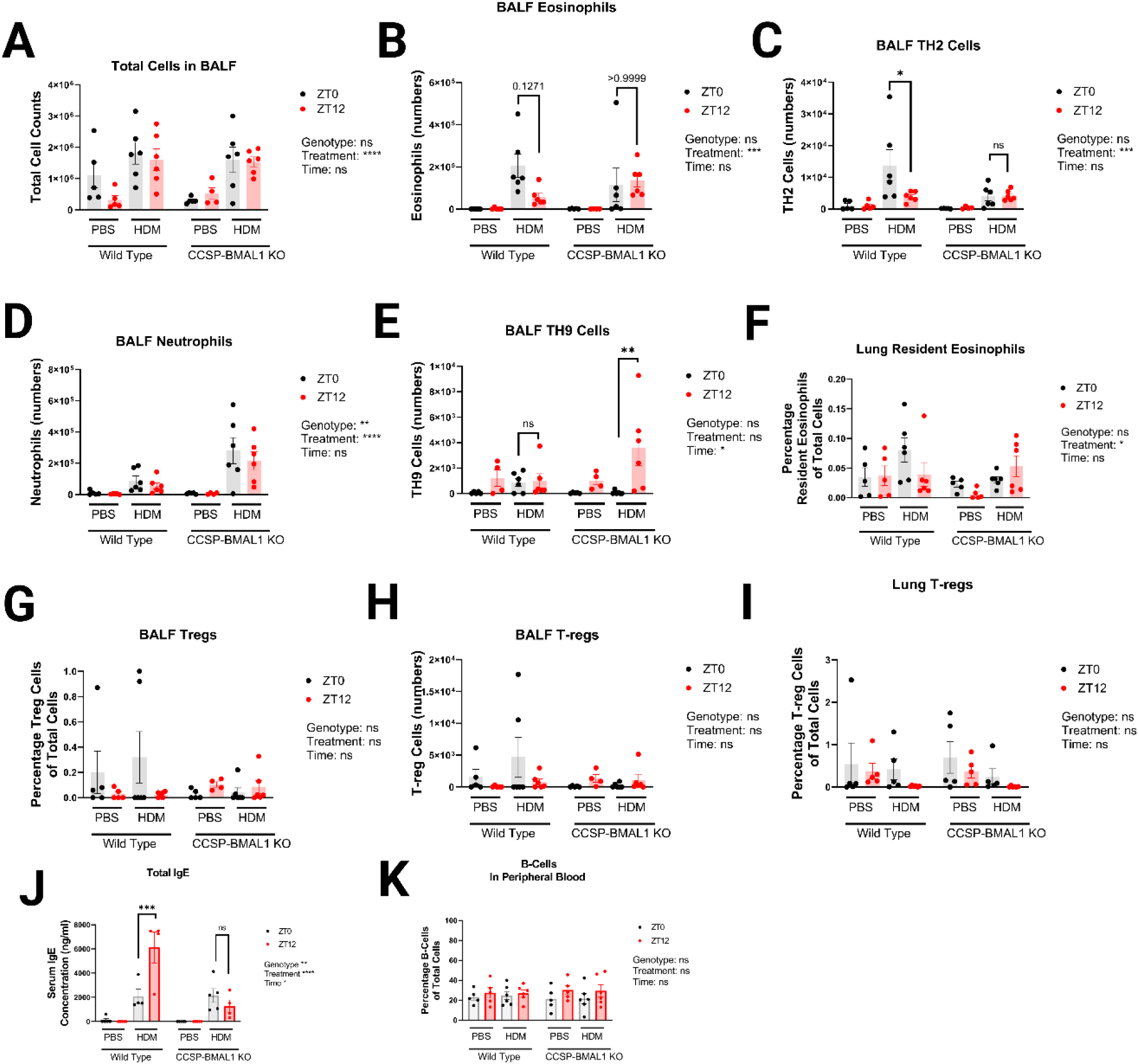
(A-I) – Immune cell subsets from HDM/PBS treated CCSP-BMAL1 KO mice and wild type littermate controls were quantified by flow cytometry from BALF, lung and spleen samples taken on day 36 of the chronic HDM exposure protocol at either ZT0 or ZT12. (A-I) – All data presented as mean +/- SEM, analysed by three-way ANOVA with post-hoc Sidak’s multiple comparison tests. N= 5 – 6, *P< 0.05, **P< 0.01, ***P< 0.001, ****P< 0.0001. (J) – Serum IgE was quantified by ELISA from peripheral blood samples taken from HDM/PBS treated mice on day 36 of the chronic HDM exposure protocol from CCSP-BMAL1 KO mice and wild type littermate controls at either ZT0 or ZT12. Data presented as mean +/- SEM, analysed by three-way ANOVA with post-hoc Sidak’s multiple comparison tests, n= 4-5, *P< 0.05, **P< 0.01, ***P< 0.001, ****P< 0.0001. (K) – Quantification of B-cells by flow cytometry in peripheral blood samples from HDM/PBS treated mice on day 36 of the chronic HDM exposure protocol from CCSP-BMAL1 KO mice and wild type littermate controls at either ZT0 or ZT12. Data presented as mean +/- SEM, analysed by three-way ANOVA with post-hoc Sidak’s multiple comparison tests, n= 5-6, *P< 0.05, **P< 0.01, ***P< 0.001, ****P< 0.0001.

**Supplementary Figure 3.**
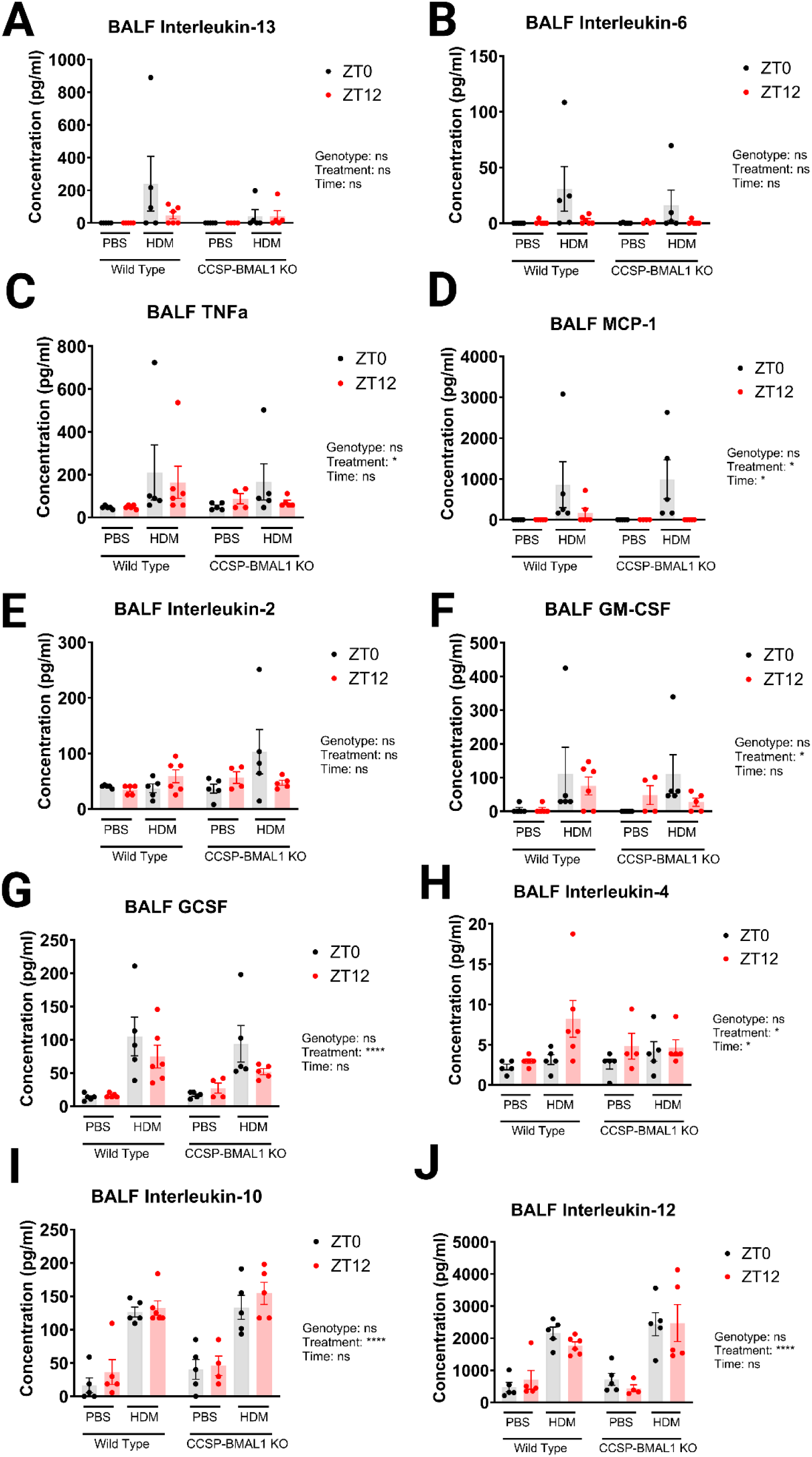
(A-J) - Quantification of cytokines present in BALF samples of HDM/PBS treated CCSP-BMAL1 KO mice and wild type littermate controls taken at ZT0 and ZT12 on day 36 of the chronic HDM exposure protocol. All data presented as mean +/- SEM analysed by three-way ANOVA with post-hoc Šidak’s multiple comparison tests. N= 5-6. *P< 0.05, **P< 0.01, ***P< 0.001, ****P< 0.0001. First level output of three-way ANOVA shown as Genotype / Time / Treatment with corresponding P value indicated.

**Supplementary Figure 4.**
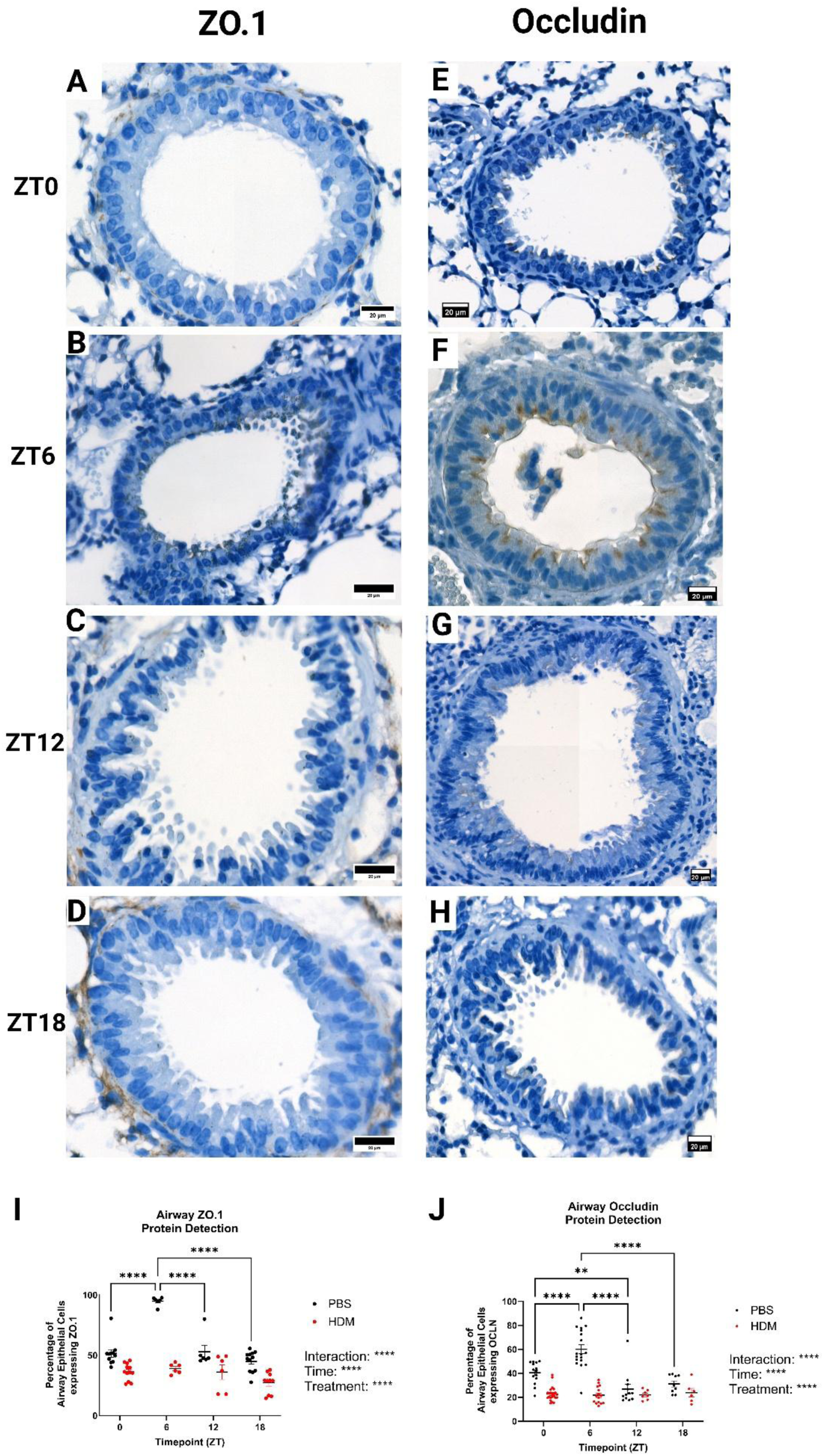
(A-D) – Representative immunohistochemical staining of airway epithelial cells in FFPE lung tissue samples stained for ZO.1 from PBS treated wild type mice at ZT0 (A), ZT6 (B), ZT12 (C), ZT18 (D). (E-H) Representative immunohistochemical staining of airway epithelial cells in FFPE lung tissue samples stained for Occludin from PBS treated wild type mice at ZT0 (E), ZT6 (F), ZT12 (G), ZT18 (H). (I) – Quantification of epithelial cells stained positive for ZO.1 across all 4 timepoints at the end of the chronic AAD protocol. N=6-12 per timepoint per treatment group, data shown as mean +/- SEM, analysed by two-way ANOVA with post-hoc Sidak’s multiple comparison tests. First level output of ANOVA shown as Interaction / Time / Treatment, *P< 0.05, **P< 0.01, ***P< 0.001, ****P< 0.0001. (J) – Quantification of epithelial cells stained positive for Occludin across all 4 timepoints at the end of the chronic AAD protocol. N=6-12 per timepoint per treatment group, data shown as mean +/- SEM, analysed by two-way ANOVA with post-hoc Sidak’s multiple comparison tests. First level output of ANOVA shown as Interaction / Time / Treatment, *P< 0.05, **P< 0.01, ***P< 0.001, ****P< 0.0001.

**Supplementary Figure 5.**
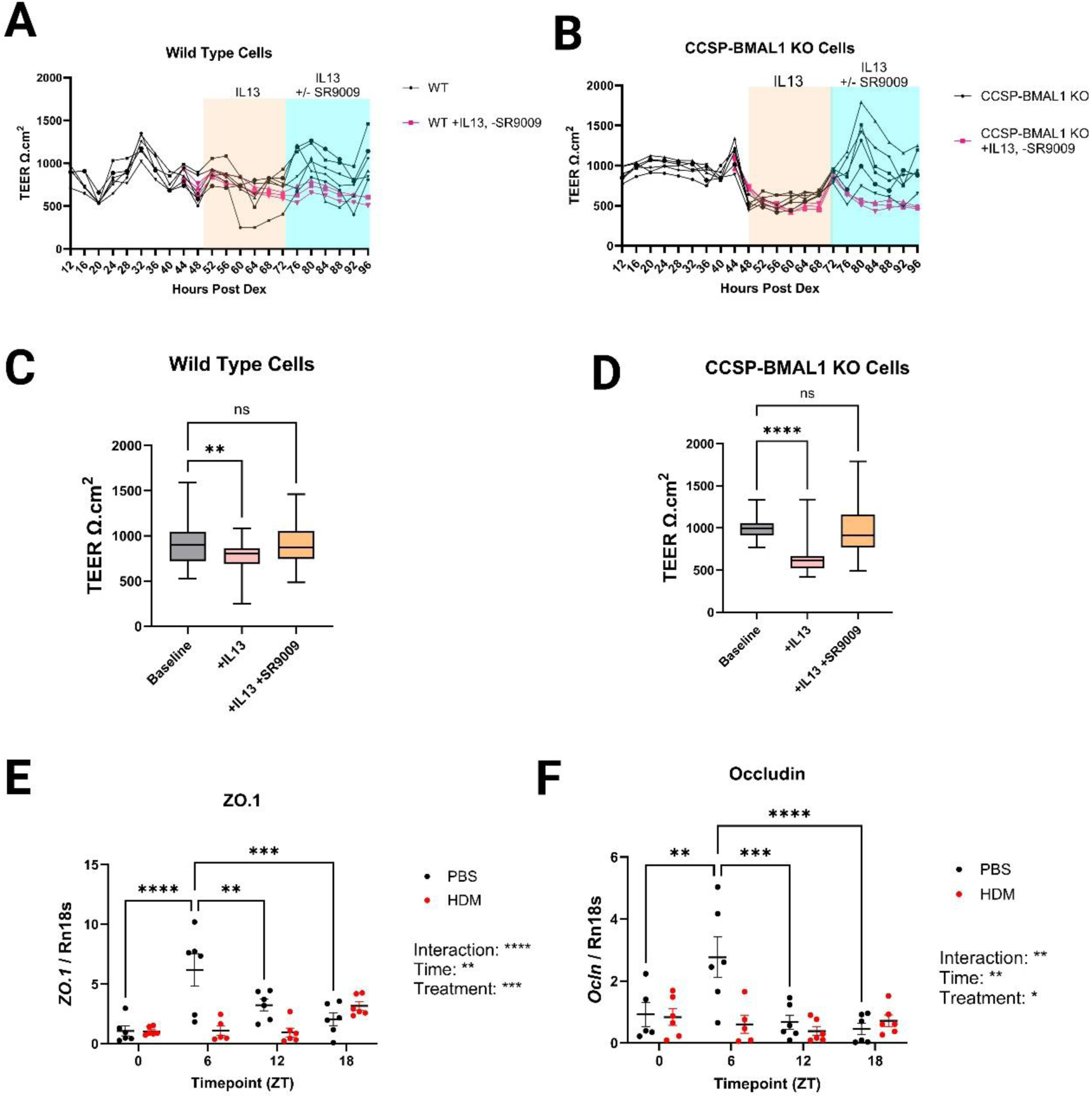
(A - B) – Trans-epithelial Electrical Resistance measurements taken from wild type (A) and CCSP-BMAL1 KO (B) air-liquid interface airway epithelial cell cultures every 4 hours from 12 hours post-synchronisation of internal clocks using dexamethasone to 96 hours post-synchronisation. At hour 48 cultures were exposed to IL13 prior to subsequently being dosed with SR9009 or DMSO as vehicle control at hour 72. N=3-6 per genotype per treatment group. (C – D) – Box and Whiskers plots showing the median and interquartile ranges of TEER recordings per each treatment group. N= 42 per treatment group, analysed by one-way ANOVA with post-hoc Dunnett’s multiple comparison tests. (E) - qPCR data showing the relative expression of tight junction gene *Zo.1* at ZT0, ZT6, ZT12, and ZT18 in whole lung samples taken from wildtype PBS/HDM-treated female mice. (F) - qPCR data showing the relative expression of tight junction gene *Occludin* at ZT0, ZT6, ZT12, and ZT18 in whole lung samples taken from wildtype PBS/HDM-treated female mice. (E-F) – qPCR data was normalised to the PBS-treated ZT0 group, and relative to the expression of housekeeping gene *Rn18s,* shown as mean +/- SEM, analysed by two-way ANOVA with post-hoc Sidak’s multiple comparison tests, n= 5-6, *P< 0.05, **P< 0.01, ***P< 0.001, ****P< 0.0001. First level output of two-way ANOVA shown as Interaction / Time / Treatment with corresponding P value indicated.

**Supplementary Figure 6.**
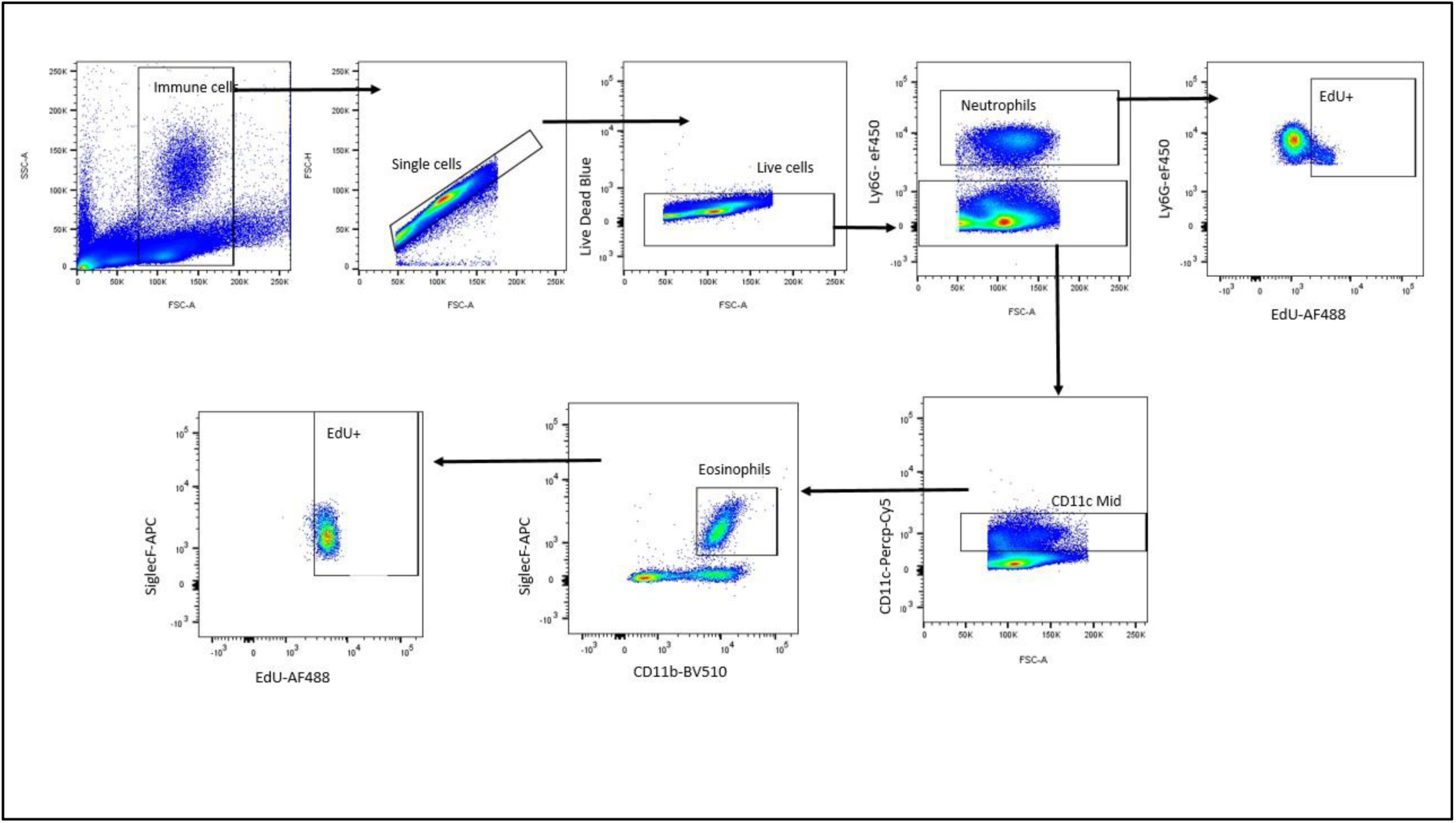
Detection of EdU by Flow Cytometry: The presence of EdU in eosinophils was detected by flow cytometry. First, cells were selected based on forward and side scatter profiles. Singlet cells were then selected, following which a Live/Dead UV stain was used to exclude dead cells. Subsequently cells were profiled in terms of LY6G expression, with LY6G+ cells being identified as neutrophils. The LY6G+ population was then profiled in terms of EdU presence indicated by the detection of the AF488 fluorophore. The LY6G-population was further analysed to identify a LYG6-CD11c (mid) population, which was subsequently analysed to identify eosinophils as a LY6G-CD11b mid SiglecF+ CD11b+ population. Like neutrophils, eosinophils were then profiled in terms of EdU presence indicated by the detection of the AF488 fluorophore.

**Supplementary Figure 7.**
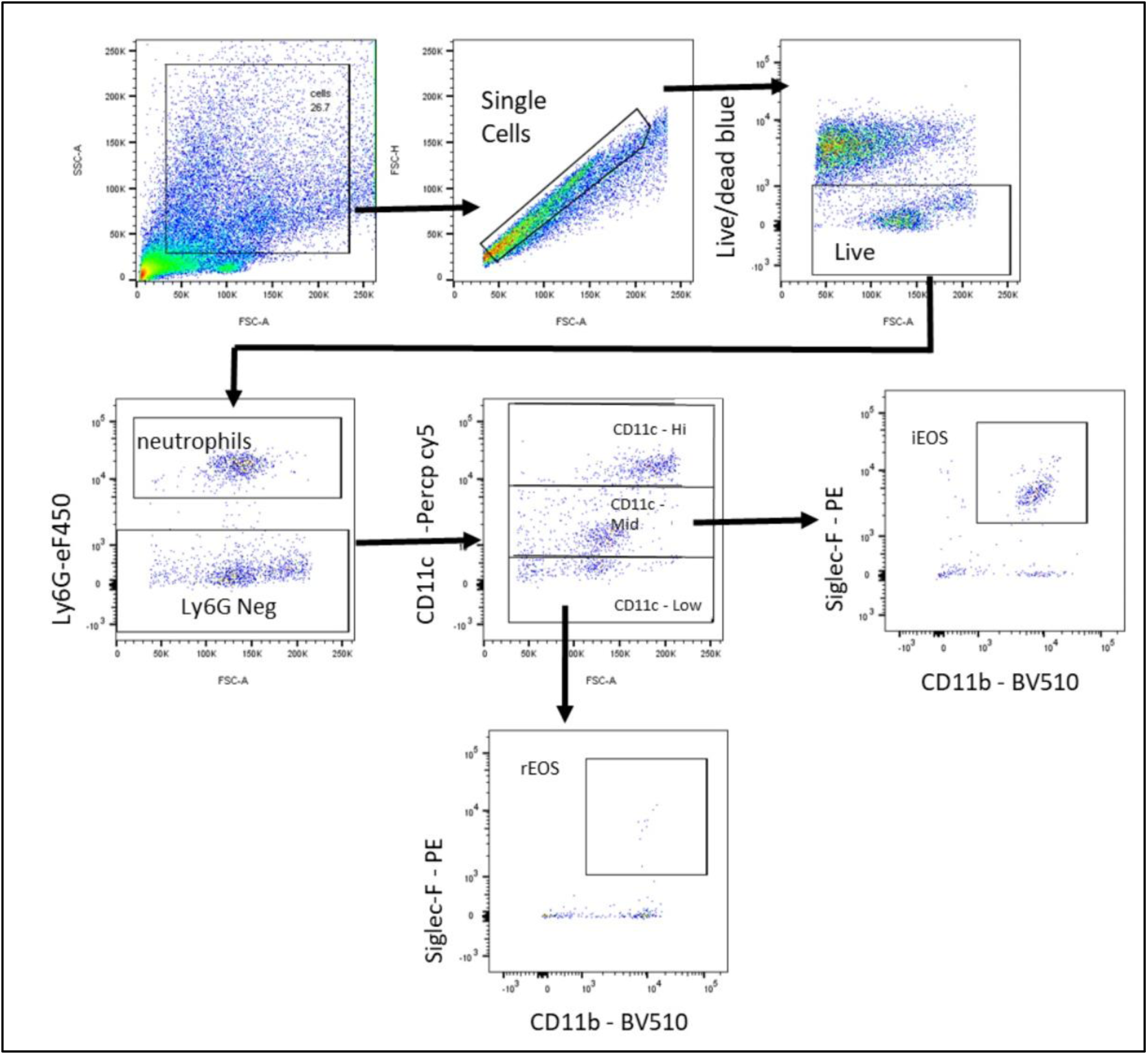
The identification of granulocytes by flow cytometry: Eosinophil and neutrophil cells were detected by flow cytometry. First, cells were selected based on forward and side scatter profiles. Singlet cells were then selected, following which a Live/Dead UV stain was used to exclude dead cells. Subsequently cells were profiled in terms of LY6G expression, with LY6G+ cells being identified as neutrophils. The LY6G-population was further analysed to identify a LYG6-CD11c (mid) population and a LY6G-CD11c (low) population. LY6G-CD11c mid cells were further analysed in terms of SiglecF and CD11b expression. LY6G-CD11c mid SiglecF+ CD11b+ cells were then identified as inflammatory eosinophils (iEos). LY6G-CD11c low cells were analysed for their expression of SiglecF and CD11b with Ly6G-CD11c low SiglecF+ CD11b+ cells being identified as resident eosinophils (rEos).

**Supplementary Figure 8.**
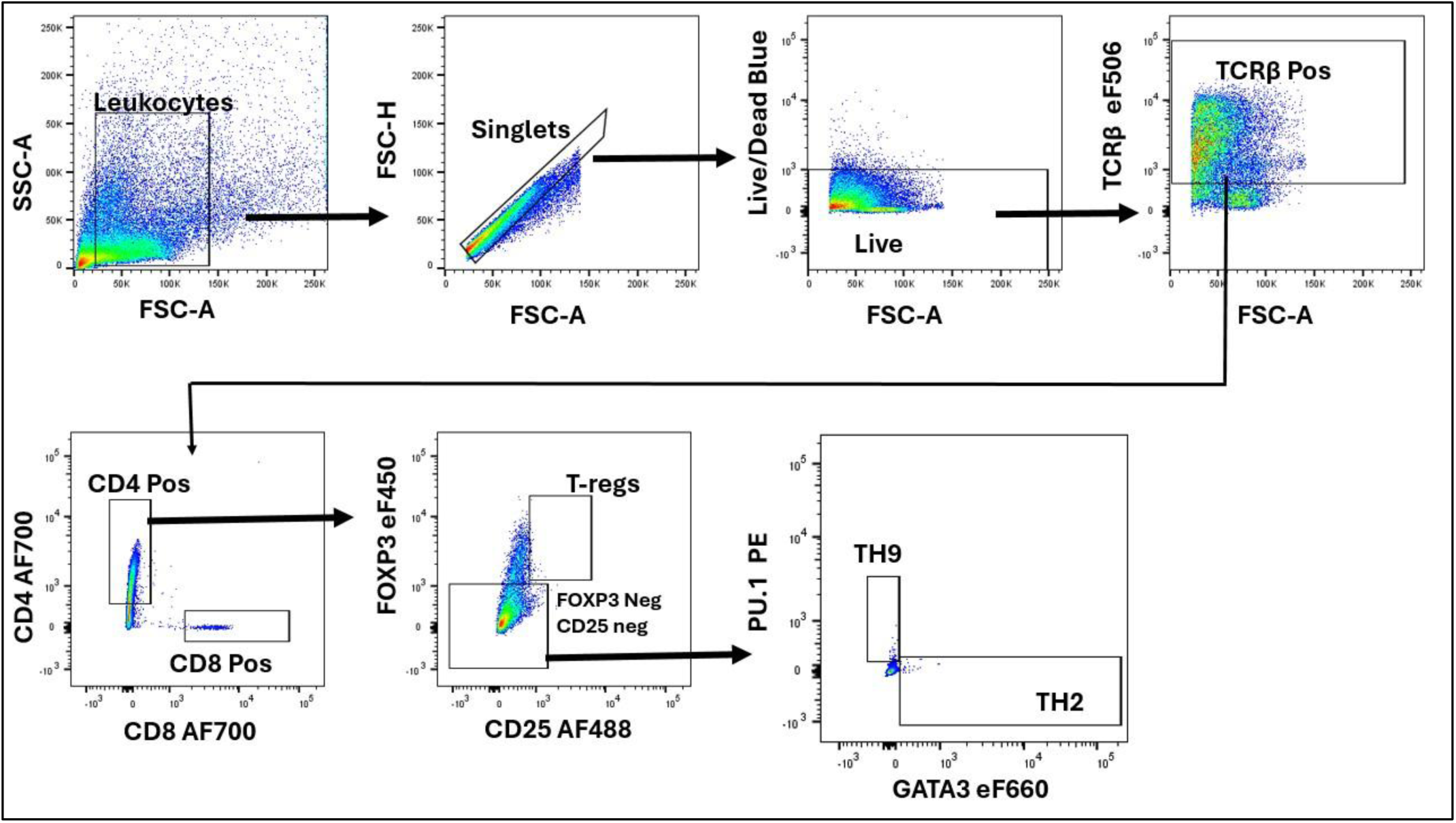
Immunophenotyping of T-cell subsets by flow cytometry: TH2, TH9 and T-reg cells were identified by flow cytometry. First, cells were selected based on forward and side scatter profiles. Singlet cells were then selected, following which a Live/Dead UV stain was used to exclude dead cells. TCRβ+ cells were then selected and taken forward to be gated CD4/CD8. TCRβ+ CD4+ cells were then analysed to show T-reg populations as TCRβ+ CD4+ CD25+ FOXP3+ cells. TCRβ+ CD4+ FOXP3-CD25-cells were then analysed to identify expression of transcription factors PU.1 (TH9 cells) and GATA3 (TH2) cells.

## Supplementary Methods

### FACS Staining

Single cell suspension samples were plated onto v-bottom 96-well plates such that each well contained 1×10^6^ cells. Plates were then centrifuged at 18’000 xg for 2 minutes and the supernatant then flicked off over a sink. Following this, cells were resuspended in each well with 50μl of CD16/32 antibody diluted 1:100 in FACS buffer. Cells were then incubated for 30 minutes at room temperature. Post-incubation 100μl FACS buffer was added to each well and the plate centrifuged to re-pellet the cells. The supernatant was discarded, and the cell pellet washed by adding 150μl of FACS buffer and repeating the centrifugation steps. Cells were resuspended in 50µl of the relevant antibody cocktail (Supplementary Table 2) containing a 1:100 dilution of antibodies for extra-cellular antigens and a 1:1000 dilution of live/dead stain (LIVE/DEAD fixable blue dead stain kit, ThermoFisher, U.S.A). Samples were incubated at room temperature for 45 minutes before being washed with 100µl of FACS buffer, centrifuged for 2 minutes at 18’000xg (4°C) to re-pellet cells. Following staining for cell surface markers, samples were fixed by resuspending cell pellets in 50µl of 4% paraformaldehyde and incubating at room temperature for 30 minutes in cases where no intracellular staining was being performed. Where intracellular staining was required, cells were fixed in 100µl of True-Nuclear fixation solution (Biolegend, USA) diluted 1:4 with the True-Nuclear fixation diluent (Biolegend, USA) and incubated for 45 minutes at room temperature. Following fixation, cell pellets were washed twice in 150µl FACS buffer before being resuspended in 150µl FCAS buffer and stored at 4°C protected from light. For intracellular antibody staining cells were permeabilised by resuspending pellets following fixation steps in 100µl of True Nuclear permeabilisation buffer (Biolegend, USA) diluted 1:10 with dH2O and incubating for 1 hour at room temperature. Cells were resuspended after permeabilisation steps in 100µl of the relevant intracellular antibody cocktail (Supplementary Table 2) containing a 1:200 dilution of antibodies. Samples were incubated at room temperature for 60 minutes before being washed twice with 150µl of Permeabilisation Buffer (Biolegend, USA). Finally, plates were centrifuged for 5 minutes at 18’000xg (4°C) to re-pellet cells, following which the supernatant was flicked off and cells were resuspended in 150µl of FACS buffer before being stored at 4°C protected from light overnight.

**Supplementary Table 1.**
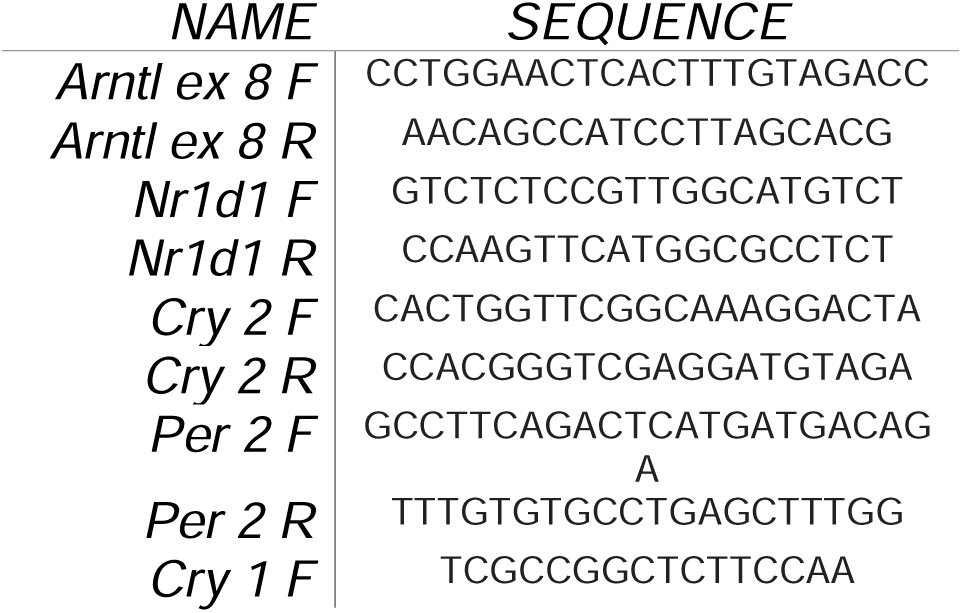
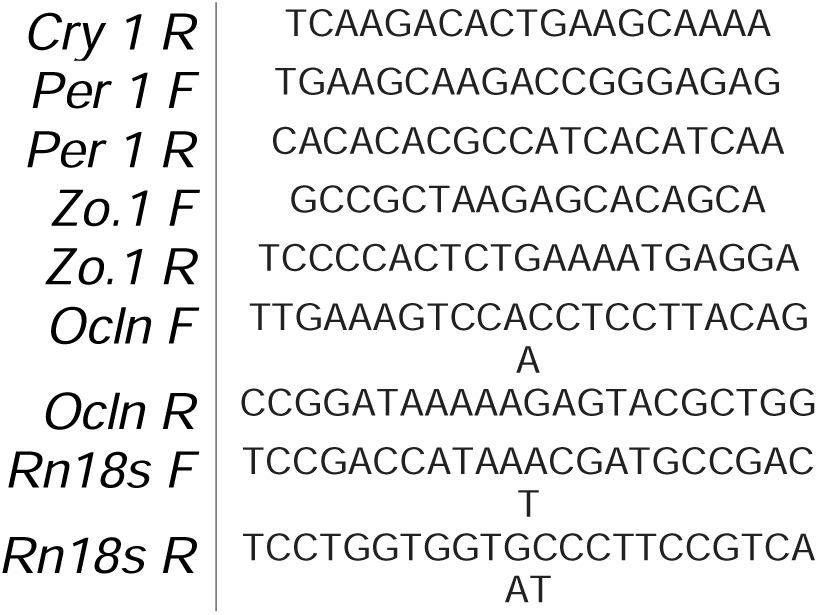

**Supplementary Table 2.**

|  | *Target* | *Fluorophore* | *Clone* | *Supplier* | *Working Concentration* |
| --- | --- | --- | --- | --- | --- |
| ***T-Helper Cell Panel*** | CD4 | AF700 | GK1.5 | Biolegend | 1:100 |
|  | CD8 | BV785 | 53-6.7 | Biolegend | 1:100 |
|  | TCRβ | eF506 | H57-597 | Invitrogen | 1:100 |
|  | CD25 | AF488 | PC61.5 | Biolegend | 1:100 |
|  | FOXP3 | eF450 | FJK-16s | Invitrogen | 1:200 |
|  | PU.1 | PE | 7C2C34 | Biolegend | 1:200 |
|  | GATA3 | eF660 | TWAJ | Invitrogen | 1:200 |
| ***BALF and Spleen Granulocyte Panel*** | CD11c | PerCP.Cy5.5 | M1/70 | Invitrogen | 1:100 |
|  | CD11b | BV510 | HL3 | BD Horizon | 1:100 |
|  | Siglec-F | PE | E50-2440 | BD Pharmingen | 1:100 |
|  | Ly6G | eF450 | 1A8-LY6G | Invitrogen | 1:100 |
| ***EdU Detection Panel*** | CD11c | PerCP.Cy5.5 | M1/70 | Invitrogen | 1:100 |
|  | CD11b | BV510 | HL3 | BD Horizon | 1:100 |
|  | Siglec-F | APC | 517007L | Biolegend | 1:100 |
|  | Ly6G | eF450 | 1A8-LY6G | Invitrogen | 1:100 |
| **Lung**<br>**Granulocyte**<br>**Panel** | CD11c | PerCP.Cy5.5 | M1/70 | Invitrogen | 1:100 |
|  | CD11b | BV510 | HL3 | BD Horizon | 1:100 |
|  | Siglec-F | PE | E50-2440 | BD<br>Pharmingen | 1:100 |
|  | Ly6G | eF450 | 1A8-LY6G | Invitrogen | 1:100 |
| **Blood**<br>**Granulocyte**<br>**Panel** | CD11b | PerCP.Cy5.5 | M1/70 | Invitrogen | 1:100 |
|  | CD11c | BV510 | HL3 | BD Horizon | 1:100 |
|  | Siglec-F | PE | E50-2440 | BD<br>Pharmingen | 1:100 |
|  | Ly6G | eF450 | 1A8-LY6G | Invitrogen | 1:100 |
|  | B220 | APC-eF780 | RA3.6B2 | Invitrogen | 1:100 |

**Supplementary Table 3.**
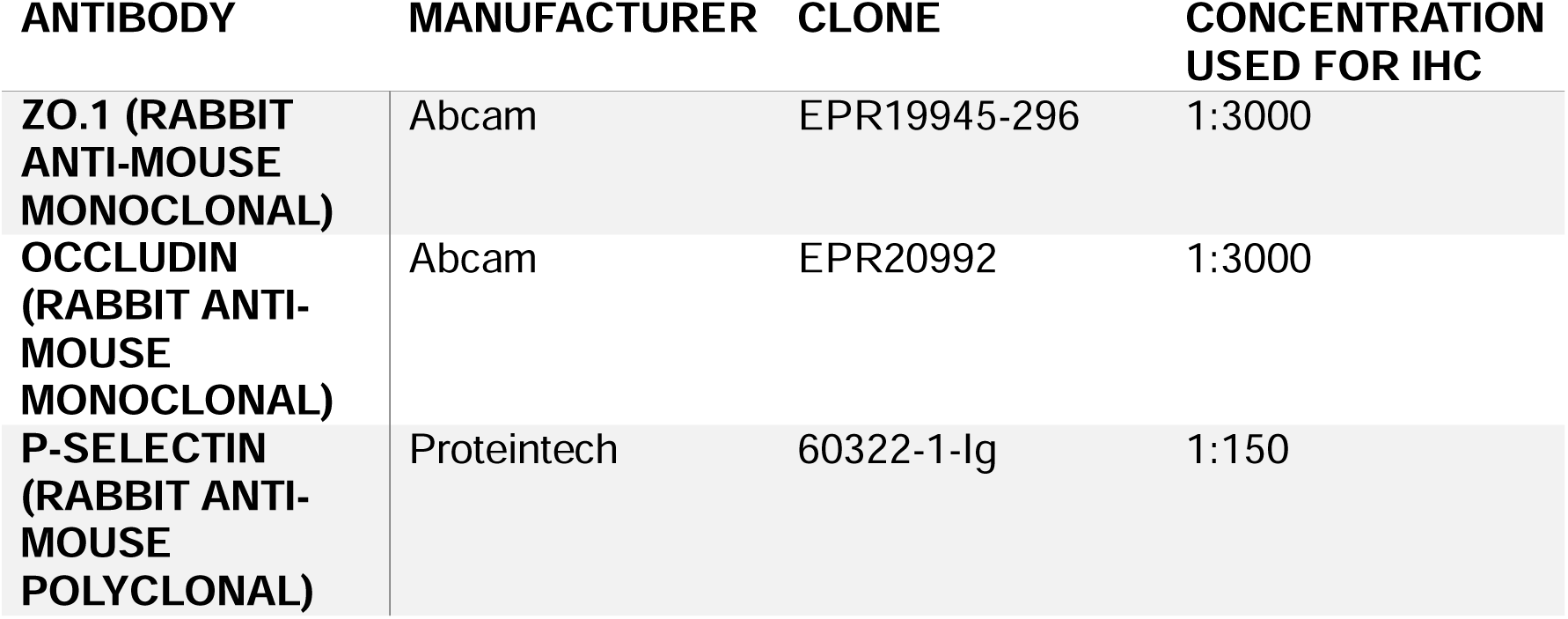

**Supplementary Table 4.**

| ANTIBODY | PRODUCT<br>CODE | COMPANY | CONCENTRATION<br>USED IN WESTERN<br>BLOT |
| --- | --- | --- | --- |
| **ZO-1 (RABBIT ANTI-MOUSE POLYCLONAL)** | 21773-1-AP | Proteintech | 1:5000 |
| **OCCLUDIN (RABBIT ANTI-MOUSE POLYCLONAL)** | 27260-1-AP | Proteintech | 1:10000 |
| **BETA TUBULIN (RABBIT ANTI-MOUSE POLYCLONAL)** | 10094-1-AP | Proteintech | 1:5000 |
| **ANTI-RABBIT IGG** | 7074 | Cell Signalling Technology | 1:2000 |

